# NPM1 mislocalization mediated by RNA Pol I inhibition alters chromatin landscape

**DOI:** 10.1101/2025.10.28.684736

**Authors:** Pierre-Olivier Estève, Sagnik Sen, Karthikeyan Raman, Ashwin Unnikrishnan, Sriharsa Pradhan

**Author notes:** These authors contributed equally (Pierre-Olivier Estève, Sagnik Sen). Correspondence: Sriharsa Pradhan, Ph.D., New England Biolabs, Inc., Ipswich, MA 01938, USA.

## Abstract

RNA polymerase I inhibition affects rRNA synthesis from rDNA clusters residing in nucleolar organizer regions (NORs). Here we have demonstrated RNA Pol I inhibition disrupts nucleolar architecture, NPM1 localization, and alters the chromatin landscape using coordinated eraser and writer enzymes. siRNA mediated depletion or dissociation of NPM1 allows HDAC1 loading on the chromatin. HDAC1 mediated deacetylation of H3K9ac creates H3K9 that undegoes SUV39H1-mediated methylation. The stripping of active histone marks leads to enrichment of repressive H3K9me3 in the genome. These altered chromatin landscape corroborates with loss of genome-wide chromatin accessibility and DNA hypermethylation mediated by DNMT1. Chromatin architectural analysis revealed disrupted nucleolar associated domains (NADs) transforming to lamin associated domains (LADs) with specific histone signatures and repressive states. The 3D nuclear architecture was remodeled by A/B compartments reorganization and loss of Hi-C loops at the H3K9ac depleted sites.

## Introduction

The ability to precisely determine the molecular effect of small molecule drugs in chemotherapy is key to future drug development and rational drug design for effective cancer treatment. CX-5461 was discovered and initially described as an RNA polymerase I (RNA Pol I) inhibitor (Khot et al., 2019), affecting rRNA synthesis from rDNA clusters residing in nucleolar organizer regions (NORs). The ribosomal RNA genes are present as multiple tandem arrays separated by intergenic spacer sequences (Sakai et al., 1995; Stults et al., 2008; Hori et al., 2021). A typical human genome generally possesses about ten NORs (Henderson et al., 1972; Stults et al., 2008). Only a subset of rRNA genes is actively transcribed. The mechanism of CX-5461 mediated inhibition was reported to act specifically on RNA Pol I by binding to SL1, thereby disrupting preinitiation complex (PIC) formation and preventing binding of RNA Pol I to the rDNA gene promoter (Drygin et al., 2011). Additional morphological studies have demonstrated the effect of CX-5461 on the mammalian nucleus using time-lapse imaging, immunofluorescence, and careful ultrastructural analysis using HeLa cervical cancer cell line (Snyers et al., 2022). Cells treated with CX-5461 displayed a profound impact on their nucleolar morphology and function including a compact, spherical-shape nucleoli with enlarged, ring-like masses of perinucleolar condensed chromatin. In addition, the authors also reported nucleolar components involved in rRNA transcription, including ribosomal DNA, RNA Pol I, and fibrillarin, to maintain their topological arrangement with reduced numbers and localization towards the nucleolar periphery. Therefore, CX-5461 not only affected nucleolar chromatin rearrangements, but it also increased the heterochromatin (Snyers et al., 2022). These morphological alterations are accompanied by a decrease in cell division leading to apoptosis. These properties have made CX-5461 as a potential therapeutic candidate for hematological malignancies (Khot et al., 2019, Ferreira et al., 2025).

Other proposed molecular mechanisms underpinning the therapeutic efficacy of CX-5461 include stabilization of G-quadruplexes (G4) and impeding topoisomerase II (TOP2) that results in activation of DNA damage response pathways leading to apoptosis (Musso et al., 2017; Xu et al., 2017; Bossaert et al., 2021; Bruno et al., 2020; Cameron et al. 2024). In a recent study, Koh et al. demonstrated CX-5461 as a synthetic lethality agent in BRCA1-/BRCA2-deficient cells, perhaps via extensive, nonselective, collateral mutagenesis in mammalian cells that exceeds known environmental carcinogens (Koh et al., 2024). Therefore, CX-5461 appears to be a small molecule chemotherapeutic reagent with more than one target in mammalian cells. Currently, only a handful of other pharmacological drugs exist that directly target mechanisms of nuclear assembly and organization, but interestingly, they can also induce cancer cell death. For example, the farnesyltransferase inhibitor R115777 inhibits the growth of B-cell lymphoma and multiple myeloma (Witzig et al., 2021). It also inhibits breast and ovarian cancer cells *in vitro* and reduces the tumor growth in xenograft model systems (Wärnberg et al., 2006).

Cancer cell sensitive to CX-5461 is also dependent on nucleophosmin (NPM1) levels. NPM1 silenced CaSki cells treated with 500 nM CX-5461 were significantly sensitized compared to control samples treated with the same dose (Ismael et al., 2019). However, the detailed mechanism and connection between NPM1 amount and CX-5461 sensitivity are not known.

NPM1 is a nucleus-cytoplasmic shuttling protein located in the nucleolus and participates in ribosome biogenesis among a host of other cellular functions (Borer et al., 1989; Grisendi et al., 2006; Frottin et al., 2019; Yu et al., 2006). NPM1 mutations are the most common genetic alteration in acute myeloid leukemia (AML), detected in about 30–35% of adult AML and more than 50% of AML with normal karyotype (Yao et al., 2023). NPM1-mutated AML is regarded as a distinct genetic entity in the World Health Organization (WHO) classification of hematopoietic malignancies (Khoury et al., 2022). Since CX-5461 disrupts the nucleolus architecture, inhibits rRNA gene expression, and is sensitive to cells with low amounts of NPM1, we hypothesized that it would have a profound impact on NPM1 binding, function, and localization in the genome. Here we have dissected the role of chemotherapeutic drug CX-5461 in nucleolus structural integrity and NPM1 localization with special emphasis on in post-translational histone modification, DNA methylation, chromatin accessibility, and gene expression. Finally, we have investigated dynamic changes in the nuclear assembly and organization in response to CX-5461, focusing on nucleolar associated domains (NADs) and lamin associated domains (LADs).

## Results

### RNA Pol I inhibition affects RNA dependent NPM1 localization, alters nucleus morphology, and structure

CX-5461 is a well-known inhibitor of RNA polymerase I (Pol I) affecting ribosome biogenesis, nucleolar morphology and function including perinucleolar condensed chromatin (Quin et al., 2016; Mars et al., 2020; Syners et al., 2022). This led us to first measure the structural changes of nucleolus and nucleus after RNA Pol I inhibition with CX-5461 on HT1080 cells. We monitored the localization of NPM1 (red) and lamin B2 (green) at the nucleolus and nucleus, respectively. Prior to CX-5461 exposure (0 hr), all NPM1 exclusively remain localized at the perinucleolus space (Fig. 1A). Within three hours of CX-5461 exposure, there was rapid loss of NPM1 from the perinucleolus space (Fig 1A). This trend continued as time progressed, and gradually the majority of NPM1 localized to the nuclear periphery (Fig. 1A, B). Quantitative pixel intensity of the NPM1 ratio between nucleus periphery and center also demonstrated gradual accumulation of NPM1 to the periphery till 12 hr of drug exposure. However, at 15 hr there was diffusion of the NPM1 signal at the nuclear periphery (Fig. 1C). Concurrent with NPM1 localization to the proximity of nuclear lamina, we also observed a gradual increase in the detachment of lamina from nuclear membrane. At 15 hr, >95% of cells displayed lamin detachment from the nuclear lamina (Fig. 1D).

**Figure 1.**
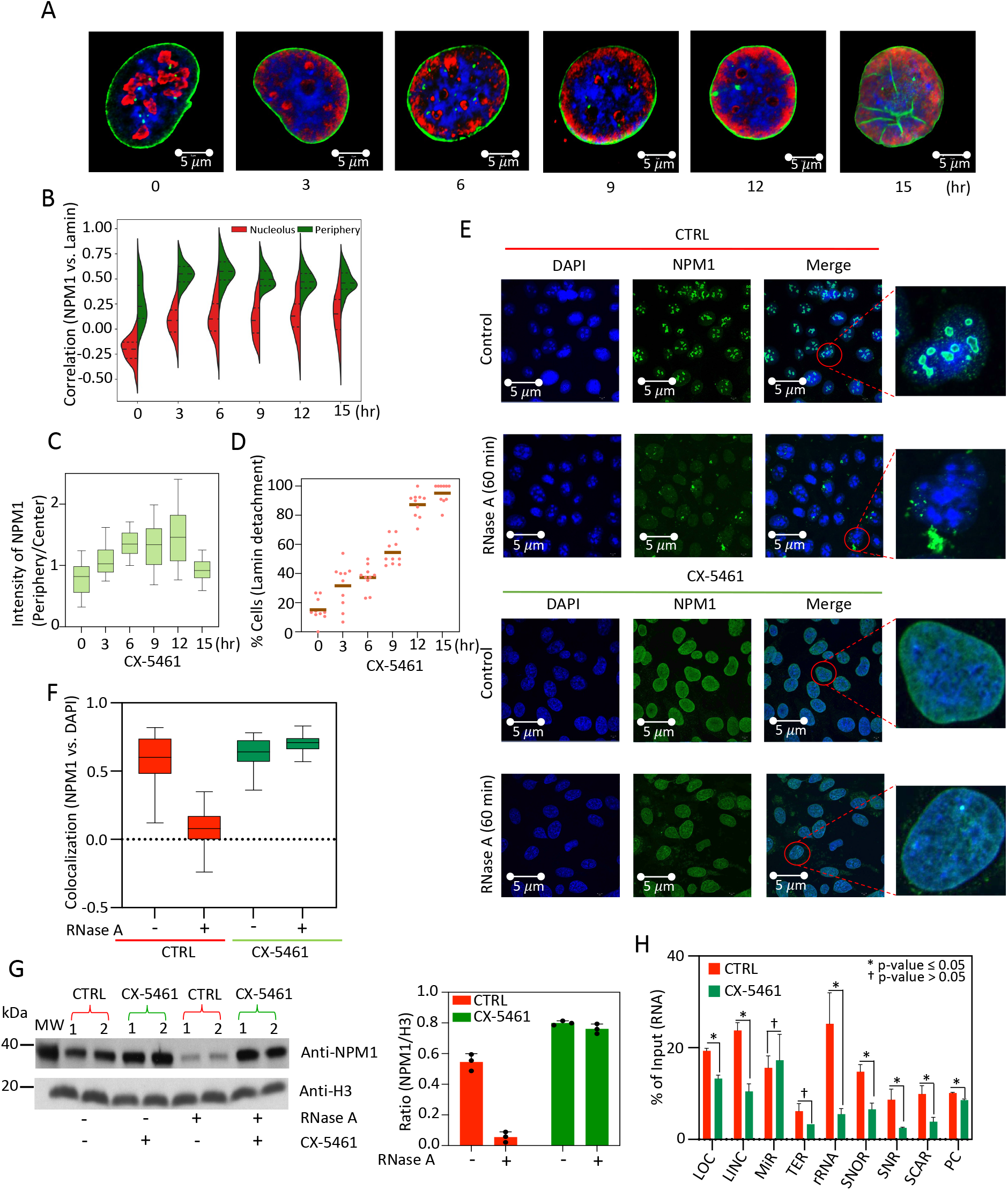
RNA Pol I inhibition by CX-5461 affects NPM1 nucleolus organization. **A**. Images showing NPM1 (red) and lamin B2 (green) localization upon CX-5461 exposure at various time interval. The nucleus is stained with DAPI. **B.** Colocalization plot of NPM1 and lamin at nucleolus and nucleus periphery based on signal intensity from microscopic images. 100 images were used for correlation calculation as shown. **C.** Ratio of NPM1 intensity at nucleus periphery vs. center, showing NPM1 diffusion. **D.** Percentage of cells showing nuclear lamina detachment with various time interval from the nuclear membrane post CX-5461 exposure. **E.** RNase A treated control and CX-5461 exposed cells displaying differential nucleolus integrity. NPM1(green) and DAPI stained nucleus are shown. **F.** Quantitative colocalization measurement using boxplot upon RNase A treatment in control and CX-5461 exposed cells. Control cells (red box) used buffer compared to RNase A treated CX-5461 cells (green box) are shown. **G.** Western blot of anti-NPM1 with control and CX-5461 exposed cell extract is shown (left panel). Each sample have two replicates and 3 biological samples tested. RNase A treated cells are indicated and the corresponding bar plot (control in red, and CX-5461 exposed in green) to demonstrated effect on NPM1, normalized to H3 control. **H.** Bar graph showing normalized percentage of NPM1 bound RNA categories, derived from RIP-seq data. Significant changes are indicated with various p values. The p-values are indicated ≤ 0.05 denotes significant losses with asterisk.

Next, we performed <u>N</u>icking <u>E</u>nzyme <u>E</u>pitope targeted <u>D</u>NA <u>seq</u>uencing (NEED-seq, Sen et al. 2025) analysis of NPM1 and control RNA Pol I to study their dynamic localization on chromatin. Indeed, we observed loss of their binding on the rDNA gene cluster post CX-5461 exposure as expected (Supp Fig. 1A). To precisely determine the dynamics of NPM1 localization, we performed relative energy analysis of NPM1 (as detailed in the Methods), using DAPI as the standard for 100 nuclei. The rate of nuclear deformation, defined as ln(k), is directly proportional to the relative energy of NPM1, whereas the relative energy of NPM1 is inversely proportional to the diffusion coefficient (DC) of NPM1, suggesting that diffusion of NPM1 is correlated to intact nuclear structure (Supp Fig. 1B). Since most of these events occurred till the first 9 hr post CX-5461 exposure, we performed LOWESS (Locally Weighted Scatterplot Smoothing) analysis between relative energy and diffusion of NPM1 and observed 6 hr to be the inflexion point for diffusion with high energy barrier (Supp Fig. 1C). We also performed a similar analysis for lamin B2 in the presence of CX-5461, for its localization in spatial context using 100 individual nuclei. We observed dynamic deposition of lamin B2 to nuclear membrane from the center till first 9 hr with a consistent nucleus size < 10 µM. However, at 15 hr the nucleus increased its size to ∼18 µM, suggesting disruption of nucleus structure and apoptosis (Supp. Fig. 1D).

Since NPM1 has RNA binding activity, we next studied the role of RNA in NPM1 localization at the perinucleolus space. We hypothesized that if ribosomal RNA is the primary determinant of NPM1 localization, then depletion of RNA or inhibition of RNA Pol I would perturb NPM1 localization. We used control and CX-5461 treated fixed HT1080 cells and incubated those cells either with RNase A or buffer as a control. Indeed, NPM1 perinucleolar colocalization was disrupted in control cells treated with RNase A (Fig 1E). On the contrary, CX-5461 treated cells displayed NPM1 throughout the nucleus due to RNA Pol I inhibition (lack of rRNA synthesis; Fig. 1E). Quantitative measurement of colocalization between NPM1 and DAPI displayed an ∼50 X reduction in control cells treated with RNase A, compared to CX-5461 treated cells, where the RNase A mediated effect was minimal (Fig. 1F). To determine if dynamic localization of NPM1 from the nucleolus to lamina following RNase A treatment promoted its diffusion out of the nucleus, we performed western blot on nuclear extract using anti-NPM1 and normalized to H3. Indeed, RNase A treated control cells lost RNA bound NPM1 from the nucleus, but not CX-5461 treated cells (Fig. 1G). Conversely, RNase H treated cells did not show any loss of NPM1 from the nucleus, in either control cells or CX-5461 treated cells (Supp. Fig. 1E), suggesting NPM1 binding is exclusively driven by RNA interactions and not through RNA-DNA hybrid. To determine the class of NPM1 bound RNA affected by CX-5461, we performed RNA Immunoprecipitation-sequencing (RIP-seq) analysis. Indeed, many different classes of RNA, including rRNA, lost NPM1 binding in CX-5461 exposed cells (Fig. 1H). Additionally, western blot analysis of total NPM1, and lamin B2 displayed no effect on their levels when normalized to beta-actin, demonstrating CX-5461 does have any effect on protein levels (Supp. Fig. 1F). As expected, RNA Pol I level decreased gradually with time. These results suggest that CX-5461 doesn’t alter the half-life of NPM1 and lamin B2 but affects their RNA binding and nuclear localization.

### Histone modification and NPM1 localization altered by RNA Pol I inhibition

Since RNA Pol I inhibition by CX-5461 affected NPM1 binding and nuclear ultrastructure, we performed Kernel Density Estimation (KDE) of NPM1 and lamin B2 bound regions for enrichment analysis at 0 and 9 hr post CX-5461 exposure. At 9 hr there was a shift of NPM1 genomic position to the right compared to 0 hr, demonstrating redistribution of NPM1 in the genome (Fig. 2A). Since NPM1 has already been shown to be an important co-activator for RNA polymerase II driven transcription and acetylation of NPM1 enhances this activity through increased histone binding and chaperone activity (Senapati et al., 2022), these led us to investigate the correlation between NPM1 binding and active histone marks (H3K4me1, H3K4me2, H3K9me3, H3K9ac, and H3K27ac) as well as repressive histone marks (H3K9me2, H3K9me3, H3K27me3, and H2AK119ub) in both control and CX-5461 treated cells using NEED-seq. (Supp. Fig. 2A). There was a high degree of correlation between NPM1 and all active histone marks in the control cells (Pearson’s coefficient, p = 0.76-0.93) but it decreased (p = 0.48-0.63) upon CX-5461 exposure (Supp. Fig. 2B). Conversely, NPM1 correlation with repressive histone marks showed a marked increase upon CX-5461 exposure (especially with H3K9me2 (p= 0.22 to p= 0.67)), a LAD associated histone mark (Supp Fig. 2C). These observations show NPM1 reduces its binding on active histone marks and increases its occupancy on repressed histone marks following CX-5461 exposure. Since we observed the migration of NPM1 to LAD regions previously (Fig. 1A, B), we compared the correlation between NPM1 and lamin B2. Indeed, the correlation between NPM1 and lamin B2 increased post CX-5461 exposure (p= 0.51 to p= 0.80), further supporting NPM1 and LAD association (Supp Fig. 2C).

**Figure 2.**
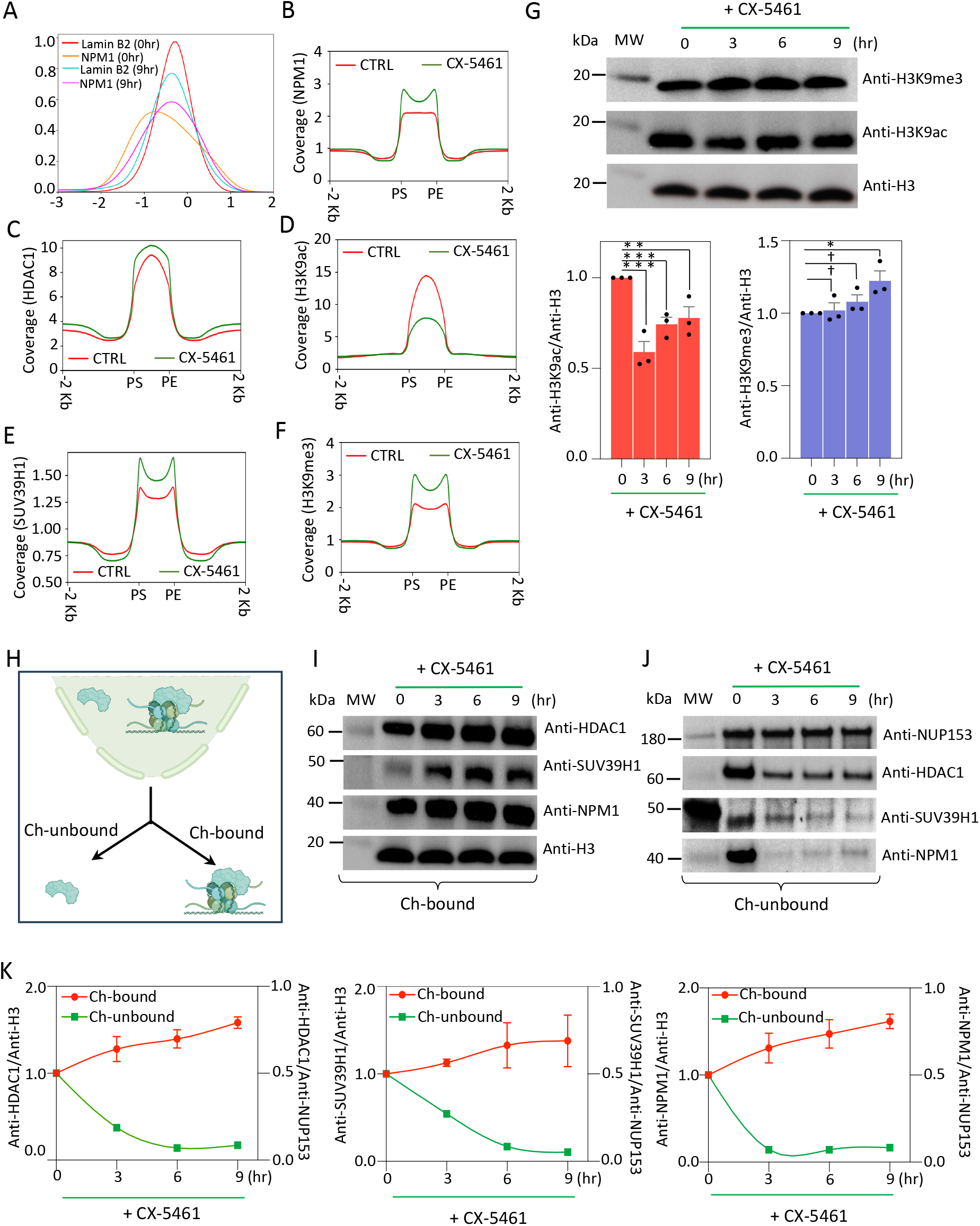
Global histone modification dynamics along with writer and eraser post RNA Pol I inhibition. **A.** Kernal Density Estimation (KDE) on NPM1 and lamin B2 coverage at 0hr and 9hr post CX-5461 incubation. The graph shows NPM1 (magenta) shifting within lamin B2 (turquoise) occupied regions upon 9hr of CX-5461 incubation, compared to NPM1 (orange) and lamin B2 (red) at 0h. **B-D**. Metagene plots showing gain of enrichment for NPM1, HDAC1 and loss of enrichment for H3K9ac. **E-F**. Metagene plots showing gain of enrichment for SUV39H1 and H3K9me3 upon CX-5461 exposure. **G.** Western blots of H3K9me3 and H3K9ac from total cell extract at various time point as indicated. Quantitative measurements of H3K9ac (red) and H3K9me3 (blue) normalized to histone H3 are shown with p-values († p-value > 0.05; * 0.01< p-value ≤ 0.05; ** 0.001< p-value ≤ 0.01; ***p-value ≤ 0.001). **H.** Schematic diagram defining the fractionation process for nuclear proteins to analyze the enrichment in chromatin bound and unbound (nucleoplasm extract). **I-J**. Western blots for chromatin bound and unbound proteins, HDAC1, SUV39H1, and NPM1 are shown. **K.** Graph showing chromatin bound fraction (red) and nucleoplasm fraction (green) for HDAC1, SUV39H1 and NPM1 at different time interval as indicated. The chromatin bound proteins were normalized to H3 and chromatin unbound proteins were normalized to NUP153.

Next, we investigated if the eraser and writer enzymes play any role in redistribution of histone marks since LADs are predominately heterochromatic in nature. We plotted enrichment plots for NPM1, H3K9ac, and eraser enzyme HDAC1, along with H3K9me3 and the writer enzyme SUV39H1 (Fig. 2B-F). There was a rapid enrichment of NPM1, SUV39H1, and HDAC1, 9 hr post CX-5461 exposure, resulting in H3K9me3 enrichment and reduction of H3K9ac. Indeed, western blot analysis of total cellular extracts post CX-5461 exposure demonstrated a global decrease in H3K9ac levels and a concurrent increase in H3K9me3 levels (Fig. 2G). In a parallel experiment, we exposed cells to BMH-21 a different RNA polymerase I inhibitor which is known to induce RNA Pol I degradation and not to activate DNA damage response (Peltonen et al. 2014). Like CX-5461 exposure, we also observed decrease in H3K9ac and a concurrent increase of H3K9me3 suggesting RNA Pol I inhibition offers common mechanism to alter chromatin modification dynamics that is independent of DNA damage response (Supp Fig. 3A). In CX-5461 exposed cells, we didn’t observe enrichment or change in protein levels for two other histone marks, H3K4me3 and H3K27me3 demonstrating mechanistic specificity of histone modification changes limited to H3K9ac and H3K9me3 (Supp Fig. 3B-C). To ensure the enriched proteins are indeed chromatin bound, we fractionated nuclear proteins to chromatin-bound, and nucleoplasm (chromatin-unbound) protein fractions post CX-5461 exposure (Fig. 2H). These fractions were western blotted and probed with antibodies. As expected, RNA Pol I level in both chromatin bound, and unbound fractions decreased (Supp Fig. 3D). Chromatin bound fraction of HDAC1, SUV39H1, and NPM1 increased with a concurrent decrease in the nucleoplasm fraction when normalized to histone H3 and the nucleoplasm marker NUP153 (Fig. 2I-K). To decipher the connection between NPM1, H3K9ac, and H3K9me3 dynamics post CX-5461 exposure, we generated KDE plots amongst them from NEED-seq profiles (Supp. Fig. 2A). Indeed, NPM1 shifted towards the right as expected, decreased in the overlap with H3K9ac and showing the loss of H3K9ac density over the CX-5461 exposure (Supp. Fig. 3E). Additionally, spreading of heterochromatic mark H3K9me3 was also observed in the same loci (Supp. Fig. 3F).

### NPM1 acts as a gatekeeper of H3K9ac and HDAC1 in cells

Since RNA Pol I inhibition increased HDAC1 occupancy on the chromatin, and additionally promoted deacetylation of H3K9ac, we analyzed the H3K9ac loss regions post CX-5461 exposure for NPM1, HDAC1, and H3K9me3 enrichment. This would allow us to investigate if there is a correlation between loss of H3K9ac and gain of H3K9me3 in the same region of the genome. We identified ∼16.4 K regions that lost H3K9ac following CX-5461 exposure (Fig. 3A). Next, we performed the coverage analysis for H3K9ac, NPM1, HDAC1, and H3K9me3 in those negatively enriched regions. As expected, following the CX-5461 exposure, these regions had very low coverage for H3K9ac. Surprisingly, NPM1 coverage was also low, supporting NPM1 and H3K9ac binding synergy (Fig. 3B; Senapati et al., 2022). However, HDAC1 and H3K9me3 were consistently enriched suggesting possible nucleosome repositioning (Fig. 3B). If indeed loss of occupancy of NPM1 leads to concurrent higher occupancy of HDAC1 and deacetylation of H3K9ac in these regions due to CX-5461 exposure, depletion of NPM1 would mimic this phenomenon and establish it as one of the gatekeepers of H3K9 acetylation. We performed an RNA knockdown experiment using three successive siNPM1 transfections along with the control siGFP in HT1080 cells, western blotted cell extracts, and measured the chromatin bound HDAC1, SUV39H1, NPM1, HP1α and control histone H3. Indeed, knockdown of NPM1 in the genome increased the chromatin occupancy of HDAC1, SUV39H1, and HP1α reflecting concurrent loss of H3K9ac and gain of H3K9me3 (Fig. 3C).

**Figure 3.**
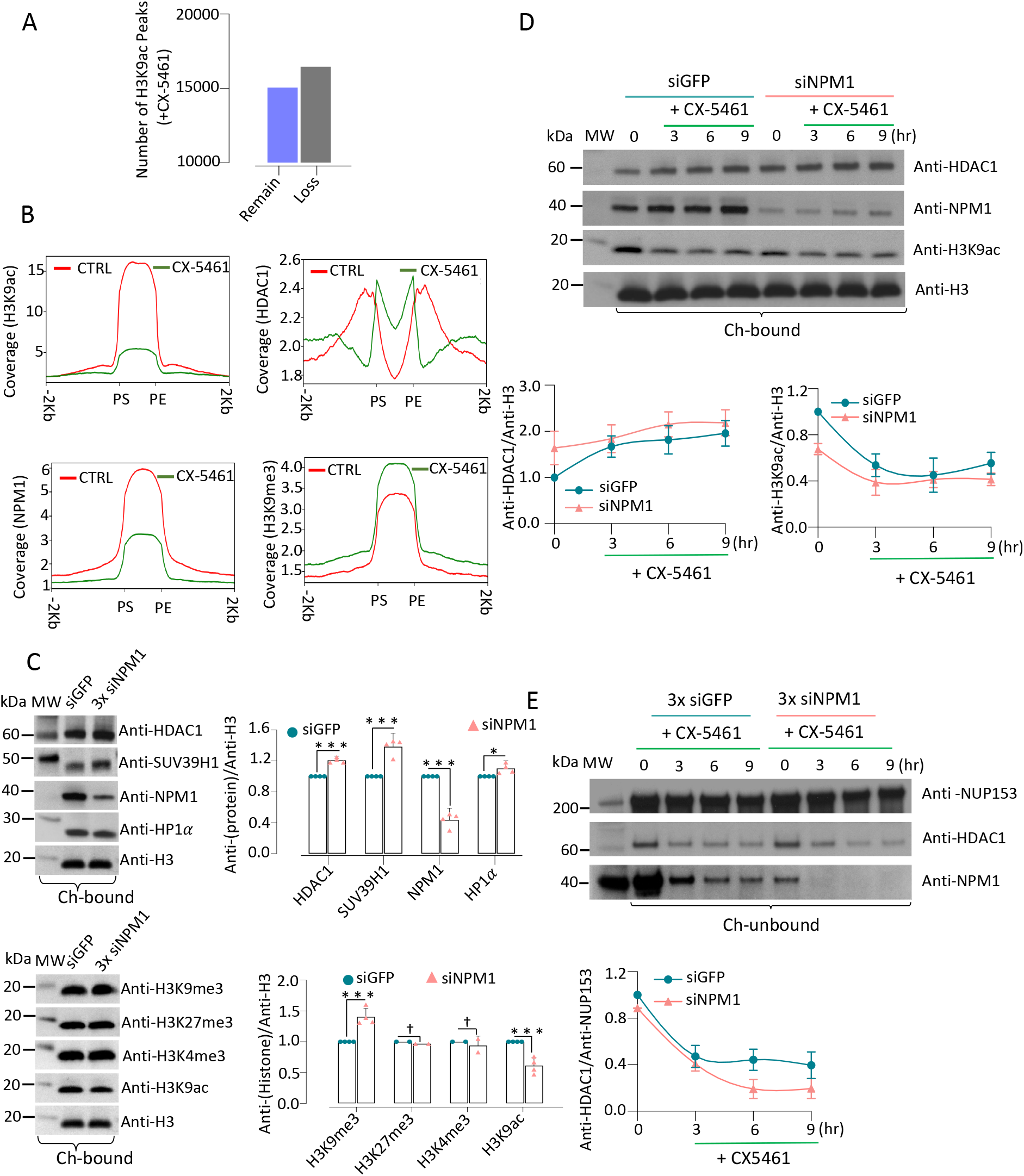
NPM1 is gatekeeper of H3K9ac deacetylation by HDAC1. **A**. H3K9ac NEED-seq peaks at control and CX-5461 exposed conditions showing ∼ 16K lost H3K9ac peaks. **B.** Metagene plots showing loss of NPM1, H3K9ac coverage within the ∼16K H3K9ac lost peaks upon CX-5461 exposure (green) compared to control (red) (left panels). However, in the same dataset HDAC1 and H3K9me3 coverage were increased upon CX-5461 exposure (green) compared to control (red; right panels). **C.** Western blots and associated quantitative bar graph following control siGFP and 3x siNPM1 (pink triangles) transfection as indicated for HDAC1, SUV39H1, HP1a in chromatin bound and unbound fraction. Histone H3 was used for normalization. Range of p-values indicated on the top († p-value > 0.05; * 0.01< p-value ≤ 0.05; ** 0.001< p-value ≤ 0.01; ***p-value ≤ 0.001) showing rate of modifications. **D.** Western blots of siGFP (green, circle) and 3X siNPM1 (pink, triangle) on chromatin bound fraction at various time point is shown. Gradual increase for HDAC1 (left panel), and gradual decrease for H3K9ac (right panel) upon 0-9hr of CX-5461 exposure are shown. The rate of modifications is normalized with histone H3. **E.** Western blots of siGFP (green, circle) and 3X siNPM1 (pink, triangle) transfection and the relative level of chromatin unbound HDAC1 is shown post CX-5461 exposure.

To decipher the role of NPM1 in HDAC1 recruitment to H3K9ac, we depleted it with 3x-siNPM1 transfection and treated the cells with CX-5461 in a time dependent manner. The control cells were treated in a similar manner with 3x siGFP (Fig. 3D, upper panel). As expected, chromatin bound fraction showed a gradual increase in chromatin occupancy for HDAC1 in siGFP cells. Enhanced binding of HDAC1 onto the chromatin was observed in siNPM1 knockdown cells. Corroborating with the HDAC1 loading on chromatin, H3K9ac levels gradually decreased on the chromatin, and it was more prominent in NPM1 knock down cells (Fig. 3D, lower panel). The chromatin unbound HDAC1 level gradually decreased due to increased loading on the chromatin upon CX-5461 exposure and was prominent in NPM1 depleted cells (Fig. 3E).

### RNA Pol I depletion mimics CX-5461 mediated inhibition

Since CX-5461 works by inhibiting RNA Pol I and recruiting NPM1 to the chromatin, we next examined if depletion of RNA Pol I would invoke a similar epigenetic response in the absence of rDNA transcription. We depleted RNA Pol I using siRNA and measured chromatin bound and unbound of HDAC1, NPM1, and H3K9me3, H3K9ac levels. We observed increased levels of HDAC1 recruitment with a resultant decrease in H3K9ac and a gain in H3K9me3 on chromatin with a concurrent decrease in the chromatin unbound fraction confirming the association between RNA Pol I inhibition and H3K9ac deacetylation (Supp Fig. 4A). Additionally, depletion of HDAC1 had no effect on H3K9me3 compared to its writer enzyme SUV39H1, where 50% reduction was observed (Supp Fig. 4B). When HDAC1 depletion was carried out in the presence of CX-5461, H3K9ac increased compared to the control (Supp Fig. 4C). These results confirm that NPM1 binding to H3K9ac acts as a barrier for deacetylation and subsequent methylation of K9 residue. In addition, either CX-5461 mediated RNA Pol I inhibition or siRNA mediated RNA Pol I depletion leads to similar epigenetic dynamics in the cell, suggesting the transcriptional state of rDNA may be a determinant in epigenetic landscape maintenance.

**Figure 4.**
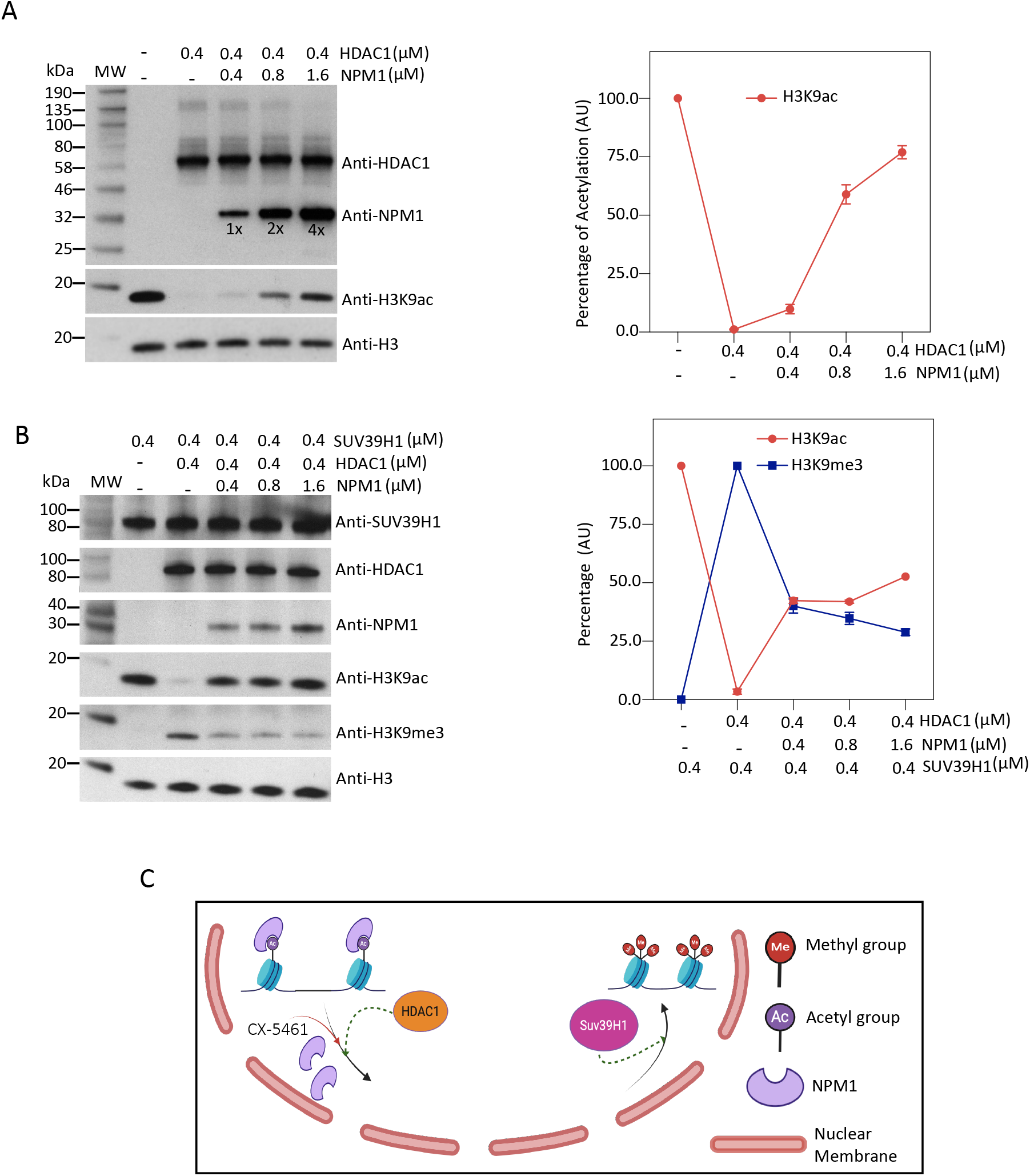
Mechanism of H3K9ac to H3K9me3 modification and role of NPM1: **A.** An in vitro incubation of HeLa total histones, HDAC1, and NPM1 to decipher role of NPM1 in HDAC1 mediated histone deacetylation. Molar concentration of NPM1 in the reaction is indicated on the top. Deacetylation reaction was monitored by western blot of H3K9ac (Histone H3 was used for normalization), and corresponding line graph on relative acetylation rate is shown (right panel). Note increased NPM1 concentration blocked deacetylation. **B.** Western blot of an *in vitro* reaction on recombinant synthetic H3K9ac, HDAC1, SUV39H1 and NPM1 to measure the conversion of acetylation to methylation (Histone H3 was used for normalization). Note complete deacetylation upon HDAC1 exposure in absence of NPM1, and gradual dose dependent increase in H3K9me3 corresponding to dose dependent increase in SUV39H1. Effect of an increase amounts of NPM1 concentration on H3K9ac to H3K9me3 conversion is shown. **C.** A schematic diagram exhibiting the possible mechanism of NPM1’s dissociation post RNA pol I inhibition by CX-5461 exposure resulting in the dynamic shift from H3K9ac to H3K9me3.

### Eraser and writer enzymes facilitate the H3K9ac to H3K9me3

We have already demonstrated that depletion of NPM1 leads to more HDAC1 loading on the chromatin, leading to reduced acetylation of H3K9ac (Fig. 3). We hypothesized, once the cells are exposed to RNA Pol I inhibitors, there would be dissociation of the NPM1-H3K9ac complex and recruitment of HDAC1. This would allow H3K9ac deacetylation by HDAC1 to H3K9, a substrate for SUV39H1 for catalytic conversion to H3K9me3. To validate our hypothesis, we perform an *in-vitro* experiment by incubating HeLa total histones with a constant concentration of HDAC1 (0.4 µM) and an increased concentration of NPM1 (0.4-1.6 µM). We western blotted and measured the quantity of H3K9ac. Indeed, HDAC1 deacetylated H3K9ac effectively, and deacetylation was blocked in the presence of increased concentrations of NPM1, confirming that NPM1 can protect H3K9ac even in the presence of HDAC1 (Fig. 4A). To demonstrate that the effect is indeed histone H3 dependent and not influenced by any other chromatin protein/(s) in the HeLa mixture, we use a purified recombinant histone H3K9ac (EPL) as substrate. As expected, 0.4 µM HDAC1 deacetylated the H3K9ac completely. Equimolar concentration (0.4 µM) of HDAC1 and SUV39H1 resulted in ∼50% of the deacetylated site trimethylated. An increased concentration of NPM1 blocked the HDAC1 mediated deacetylation, demonstrating the loss of acetylation at K9 leads to its trimethylation (Fig. 4B). Therefore, sequential removal of the NPM1 reader, and recruitment of epigenetic erasers and writers on the chromatin facilitate a dynamic shift in chromatin structure, perhaps due to deacetylation coupled with methylation of H3K9 (Fig. 4C).

### RNA Pol I inhibition by CX-5461 decreases global chromatin accessibility

Since CX-5461 participates in facilitating the H3K9ac to H3K9me3 switch on chromatin, we hypothesized that it would have a bearing on chromatin accessibility. Therefore, we performed NicE-seq (Ponnaluri et al. 2017) on both control and CX-5461 treated cells. Indeed, the chromatin accessibility on CX-5461 treated cells was reduced (Fig. 5A, B). When we examined the transcription start sites (TSS) and enhancer start and end (ES, EE) positions, the degree of decrease in chromatin accessibility was much more pronounced (Fig. 5C). Amongst these accessible peaks, ∼19k peaks were differentially modulated, with vast majority (∼97%, or ∼18k) were negatively enriched (Fig. 5D). When we analyzed these negatively enriched peaks, we found loss of NPM1 and H3K9ac coverage compared to the control (Fig. 5E, F). In contrast, increases of H3K9me3 coverage were observed in the same loci of CX-5461 treated cells, demonstrating chromatin regions with dynamic shift of histone marks associated with accessibility (Fig. 5G). This loss of chromatin accessibility and enrichment of H3K9me3 led us to investigate if there is a change in DNA methylation in these regions, a primary signature of gene silencing. We performed EM-seq in both control and CX-5461 treated genomic DNA and analyzed per CpG DNA methylation change, shown in violin plots, since they allow comparison of distribution of different groups. Indeed, the negatively enriched region of accessible chromatin showed a gain in DNA methylation (Fig. 5H). Similarly, peaks representing H3K9ac and H3K9me3 showed DNA methylation gain, with a larger skew of hypermethylation in H3K9me3 peaks (Fig. 5I). Amongst the regions of the genome that showed a switch from H3K9ac to H3K9me3, there was an increase in DNA methylation, suggesting a repressive chromatin state is associated with DNA methylation (Fig. 5J). The increase in DNA methylation was further corroborated by increased loading of DNMT1 but not DNMT3A and B to the chromatin, confirming the role of aberrant DNA methylation by DNMT1 (Supp Fig. 5).

**Figure 5.**
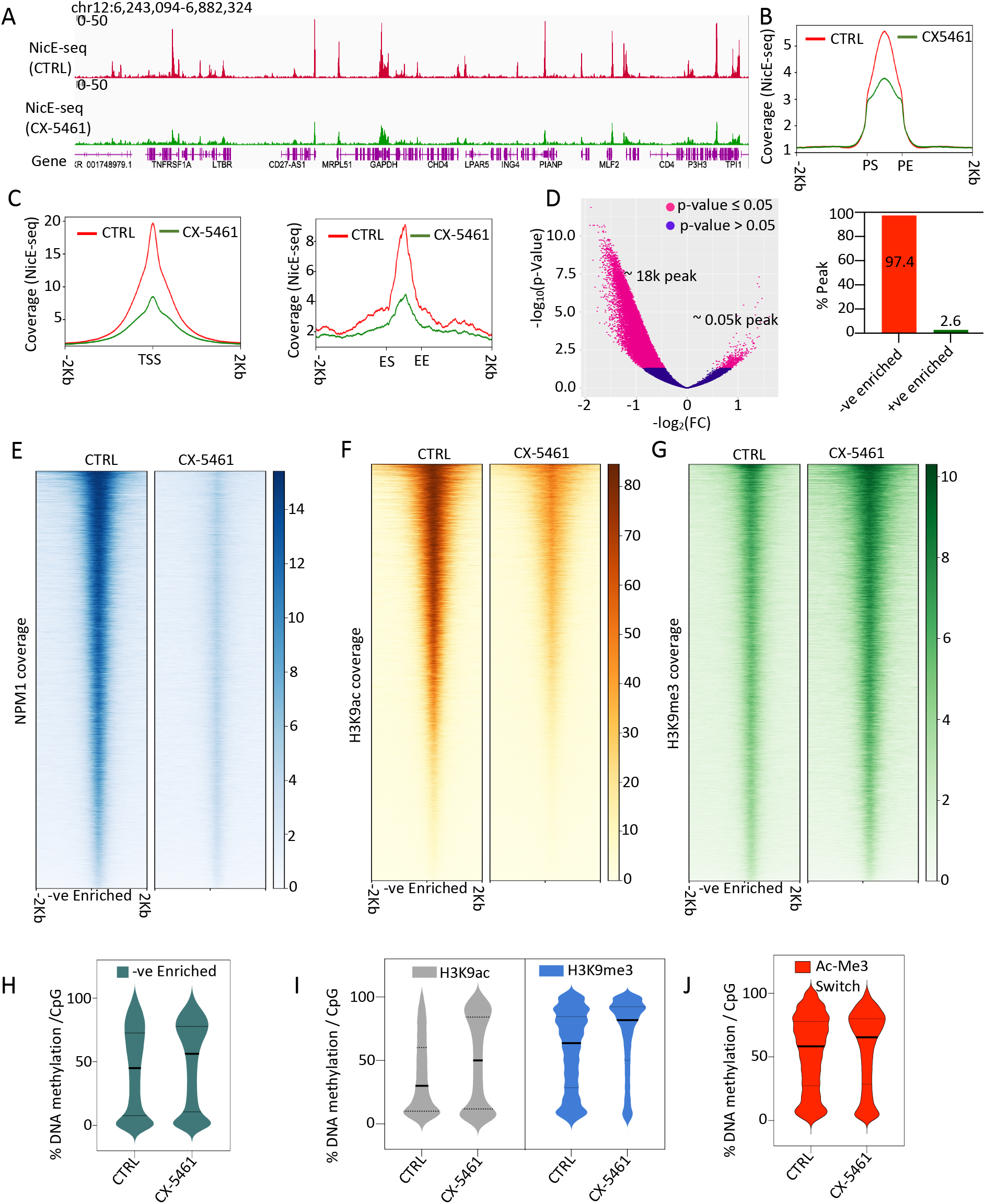
Chromatin accessibility is altered by RNA Pol I inhibition by CX-5461. **A.** IGV screenshots showing loss of NicE-seq signals between control and CX-5461 treated cells. **B.** Metagene plots showing the loss of chromatin accessibility within NicE-seq peaks, PS- peak start, and PE- peak end indicated at the bottom. **C.** Metagene plots for TSS (based on hg38) regions, and enhancer regions are shown. ES and EE indicates start and end of enhancer elements. **D.** Volcano plot and associated bar graph from DiffBind output showing significant loss of peaks (∼18K, about 97.4% of total differential peaks). **E-G.** Heatmaps on negatively enriched NicE-seq peaks showing loss of enrichment for NPM1 and H3K9ac, whereas gain of enrichment for H3K9me3. **H-J**. Violin plots exhibit changes in DNA methylation within negatively enriched NicE-peaks (sea green), H3K9ac NEED-seq peaks (grey), H3K9me3 NEED-seq peaks (sky-blue), and H3K9ac peaks that got converted to H3K9me3 peaks (Ac-Me3; red) upon CX-5461 exposure, displaying a trend for DNA hypermethylation.

### Chromatin reorganization post CX-5461 exposure

Distribution of nucleolus NPM1 to lamin and its interaction with chromatin at the nuclear periphery led us to investigate if NAD (nucleolar associated domain) and LAD (lamin associated domain) are affected in the nucleus. We analyzed H3K9me3, H3K27me3, and lamin B2 to decipher the epitope bound DNA regions, particularly NADs (using NADfinder), in both control and CX-5461 exposed cells. NADs are characterized as regions of repressive low gene density, repeated DNA sequences, and repressed chromatin states exhibiting H3K9me3 for type I NAD (NAD I) and H3K27me3 for the type II NAD (NAD II) subclass. Systematic analysis of H3K9me3 and H3K27me3 modifications at 20 kb bin size domain calling resulted in an additional 9.5 K NAD I regions (8.5 to 19.9 K) and a small but significant 2.6 K loss in NAD II regions (8 to 5.3 K) post CX-5461 exposure. Similar analysis for lamin B2 bound regions displayed significant increase of lamin B2 occupancy domains (44 K) following CX-5461 exposure (Fig. 6A). A new pattern of distribution of NAD I, NAD II, and LADs was observed in IGV suggesting that RNA Pol I inhibition affects the genome organization (Fig. 6B). This observation led us to hypothesize that NADs may be dynamically linked with LADs, and RNA Pol I inhibition plays a central role in redistribution of NADs. Since NADs and LADS are interlinked in the human genome, we calculated the overlapping percentage of NAD I and II on LADs between control and CX-5461 treated cells. Indeed, there was 35% and 50% increase in overlap with LADs for NAD I and NAD II, respectively, in CX-5461 exposed cells (Fig. 6C). Next, we studied the distribution dynamics of two nucleolus bound protein markers, nucleolin and NPM1, in control and CX-5461 exposed cells. NPM1 and nucleolin bound domains were used in NADfinder for NAD identification and designation. Since both NAD I (H3K9me3 marks) and NAD II (H3K27me3 marks) define constitutive and facultative heterochromatin, respectively, we examined if RNA Pol I inhibition has a differential impact on either of the subclasses. We derived the NADs associated with NPM1 or nucleolin and divided them into various subtypes (NAD I, NAD II, and both NAD I-NAD II overlapping domains (NAD I-II)) in both control and CX-5461 treated cells. The number of NAD II, principally associated with facultative heterochromatin decreased in number, concurrent with an increase in NAD I that is associated with constitutive heterochromatin after CX-5461 exposure (Fig. 6D). In the control cells most NPM1 bound NADs remained dissociated from LADs. However, in the CX-5461 exposed cells, these regions shifted to LADs (Fig. 6E). Enrichment plots for NPM1 bound LAD domain were higher after CX-5461 treatment supporting this observation (Fig. 6F). A similar shift of NAD to LAD was observed for nucleolin bound chromatin and its enrichment (Fig. 6G, H).

**Figure 6.**
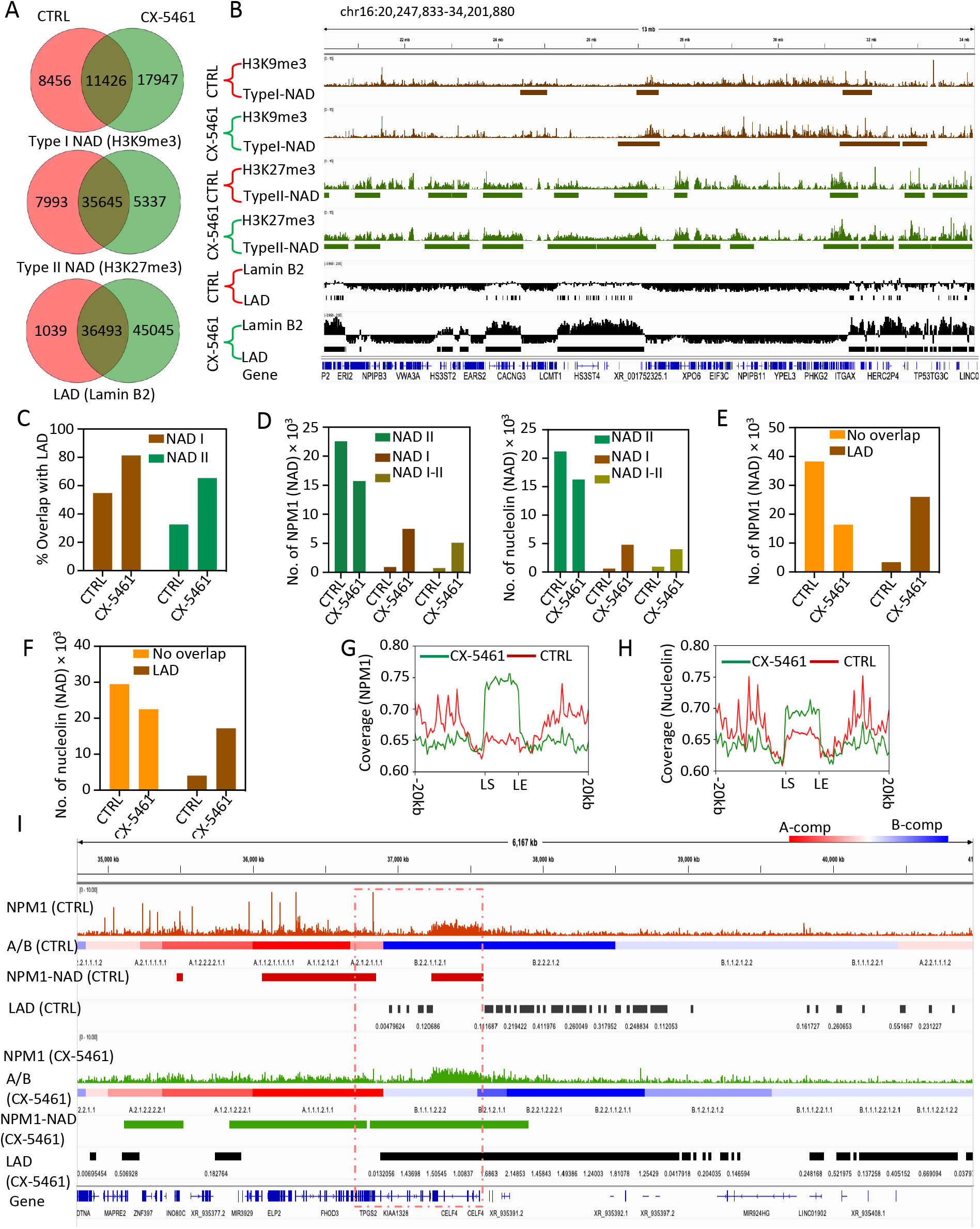
Re-distribution of chromatin domains following RNA Pol I inhibition. **A.** Venn diagrams of NAD I (H3K9me3 signature), NAD II (H3K27me3 signature), and LADs between control and CX-5461 exposed cells are shown. Changes in numbers of domains are indicated (∼9.5 K increase, ∼2.6 K decrease, and ∼44 K increase) following RNA Pol I inhibition by CX-5461. **B.** IGV screenshots of NEED-seq signals and corresponding domain bed regions are shown. NADs and representative histone marks (H3K9me3 and H3K27me3) along with the LAD domains demonstrating its expansion are shown. **C.** Bar graphs showing an increase of NAD I (brown, 35%) and NAD II (green, 50%) with LAD overlapping regions. **D.** Bar graphs showing the dynamic distribution of NPM1 and nucleolin based NAD within NAD subtypes (NAD I (brown), NAD II (green), and both NAD I-NAD II overlapping domains (NAD I-II, yellow)) in both control and CX-5461 treated cells. It shows a significant decrease in NAD II occupancy and increases in NAD I occupancy. **E.** Bar graph showing a significant increase of NPM1 based NADs within LADs. **G.** Similar to the NPM1 trend in E and showing increase in the nucleolin based NADs within LAD region in CX-5461 treated cells compared to control cells. **F-H.** Metagene plots showing NPM1 and nucleolin enrichment within LAD regions (± 20 Kb) in CX-5461 exposed (green), and control cells (red). **I.** Comparative IGV screenshot displaying NPM1 localization in A-compartment (range of red) and B-compartment (range of blue). NPM1 based NADs are expanded within LADs regions, followed a systematic shift of A-compartment to B-compartment (red dotted box) upon RNA Pol I inhibition.

In addition, both NPM1 and nucleolin NAD I-II also increased in numbers in response to RNA Pol I inhibition. This demonstrates the dynamic switch from facultative to constitutive heterochromatin for gene silencing. We further validated the heterochromatin switching phenomenon by performing global chromatin state transition for NAD I, NAD II, and LADs in control and RNA Pol I inhibited genomes. Indeed, post-RNA Pol I inhibition, NAD I representing constitutive heterochromatin, was more prominent in TSS and genic enhancers; NAD II was lost at 5’ and 3’ regions of genes along with genes associated with strong transcription. However, LAD gain was most prominent in transcriptionally active regions of the genes, particularly on enhancer elements (Supp Fig. 6A). Taken together, these results prove that RNA Pol I inhibition alters the structural integrity of the mammalian nucleus by altering transcriptional output of genes and chromatin state.

### RNA Pol I inhibition alters chromatin architecture

To understand the dynamic of chromatin architectural changes following RNA Pol I inhibition, we compared lamin and NPM1 in chromosomal conformational context using a multiomic paradigm based on lamin and NPM1 NEED-seq and Hi-C. We generated 10 subcompartments for both lamin and NPM1 by mapping the lamin or NPM1 in principal component 1 (PC1) of Hi-C in 25 kb bins. These subcompartments were compared for colocalization based on the Z-score normalized Jaccard coefficient. We observed significant enrichment of positive Z-scores between lamin and NPM1 for cells exposed to CX-5461 compared to the control (Supp Fig. 6B). The transitions in the Z-score (-ve to +ve) confirms NPM1 relocalization to LAD regions where each cluster mimics localization in chromosome conformation context. In addition, we also examined the effect of CX-5461 on topologically associated domains (TADs) in the genome that are closely associated with LADs and their influence on overall genome architecture. Indeed, the number of TADs did not change under CX-5461 exposure and remained at ∼6 K, but their size increased (Supp Fig. 6C, D). Next, we identified TADs and overlapping TADs with LADs in control vs. CX-5461 exposed cells (Supp Fig. 6E). We observed an increase of overlapping TADs with LADs and proportional loss of TADs. Additionally, Hi-C plots of chromosomes (ex: chr 21 with rDNA loci) confirm expansion of LADs within interactions (Supp Fig. 7A). When we overlap the NPM1 localization in the genome with A/B compartment architecture and positioning of corresponding LADs between control and CX-5461 exposed cells, we observed expanded NPM1-NADs to LADs. However, the expansion of B compartments was at the expense of A-compartment losses (Fig. 6I). This was further supported by NPM1 enrichment in the subtype of B-compartments compared to the subtype of A-compartments (Supp Fig. 7B). Additional studies on the differentially modified loops demonstrated that ∼60% of the loops were lost after CX-5461 exposure containing enhancer-enhancer, promoter-promoter, and promoter-enhancer interactions. These results supported loss of chromatin accessibility (Supp Fig. 8A, B). This was further validated by enrichment analysis at loop anchors corresponding to the loss of enrichment for NPM1 and H3K9ac and gain of H3K9me3 enrichment upon CX-5461 exposure (Supp Fig. 8C). In summary, chromosomal conformation analysis confirmed the spreading of constitutive heterochromatic regions propagated by NPM1 re-localization in the mammalian cells.

## Discussion

Transcription of the ribosomal RNA genes (rDNA) in the nucleolus, which encodes the three largest ribosomal RNAs (rRNA), is mediated by RNA polymerase I. This is a key regulatory step for ribosomal biogenesis and gene expression. Clinically, an increase in rRNA expression plays an active role in cancer cell proliferative capacity via control of cellular checkpoints and chromatin structure. Therefore, inhibition of RNA Pol I has gained interest in the cancer therapeutic research. CX-5461 is one of the small molecule inhibitors of RNA Pol I. It functions in part by inhibiting the formation of the PIC onto the rDNA template by specifically interfering with the interaction between SL-1 and RNA Pol I (Whiten et al., 2008). Another mechanism of CX-5461 is to promote differentiation in a MYC-interacting zinc-finger protein 1 (MIZ1)- and retinoblastoma protein (Rb)-dependent manner (Otto et al., 2022). In addition, CX-5461 is also demonstrated to increase the perinucleolar condensed chromatin, particularly the formation of a distinct DAPI-positive ring around nucleoli. The perinucleolar compartment is known to be transcriptionally repressive in nature and is enriched in silenced chromatin NAD (nucleolus-associated domain) with two repressive histone modifications (H3K9me2 and H3K27me3; Snyers et al., 2022). These observations suggest CX-5461 can affect gene expression, epigenomic dynamics, and chromatin structure in mammalian cells.

Here we showed that RNA is a bridge polymer between NPM1 and heterochromatin. A variety of RNA remains bound with NPM1, although the major class is rRNA, perhaps partly due to its abundance. This observation adds to the functional role of NPM1 in processing and transportation of multiple different RNAs, including regulating RNA splicing, polyadenylation, mRNA stability, mRNA localization, and translation. Indeed, mutations in the RNA binding domain of the NPM1 gene, specifically within exon 12, are strongly associated with acute myeloid leukemia (AML; Zarka et al., 2020). Specifically, we demonstrated NPM1 binds to NADs at the periphery of the nucleoli, and CX-5461 disrupts their association and facilitates redistribution of NPM1. In a recent study, part of the RNA-binding surface of EZH2 is required for chromatin modification at H3K27, yet this activity is independent of RNA. By careful structural studies, the RNA-binding surface within EZH2 was shown to be essential for chromatin modification in vitro and in cells, through interactions with nucleosomal DNA (Gail et al., 2024). Such a scenario for NPM1 may exist where the mutation of the RNA binding domain or blocking the binding activity may impact the epigenetic landscape change during development and cancer.

In an early immunocytochemistry study in normal human fibroblast and cancer cells, NPM1 knock down altered the distribution of H3K27me3 and H3K9me3 heterochromatin around nucleoli (Olausson et al., 2014). However, in our study NPM1 knock down in HT1080 cells resulted in higher loading of HDAC1, SUV39H1, and HP1 alpha with a concurrent decrease in H3K9ac and an increase in H3K9me3 on the chromatin. The same phenomenon was also observed if the cells were treated with CX-5461, confirming NPM1 acts as a central node for maintaining epigenetic landscape. This observation is particularly novel since CX-5461 has not been shown to be an epigenetic modulator. Over the past two decades, a growing array of small molecule drugs targeting epigenetic enzymes such as DNA methyltransferase, histone deacetylase, isocitrate dehydrogenase, and enhancer of zeste homolog 2 have been thoroughly investigated and implemented as therapeutic options, particularly in oncology, and many are currently in clinical trials. These epigenetic inhibitors modulate epigenetic landscape, like RNA Pol I Inhibitor CX-5461. Mirroring epigenetic inhibitors, we have previously shown that disruption of DNA methyltransferase activity also remodels genome compartmentalization whereby some domains lose H3K9me3-HP1α/β binding and acquire the neutrally interacting state while retaining late replication timing. In addition, H3K9me3-HP1α/β heterochromatin is permissive to loop extrusion by cohesin but refractory to CTCF binding, confirming a dynamic structural and organizational diversity of the silent portion of the genome and between the chromatin state and chromosome organization mediated by DNA methylation and the 3D genome (Spracklin et al., 2023).

Our results show that CX-5461 mediated RNA Pol I inhibition leads to loss of chromatin accessibility with concurrent gain of H3K9me3, DNA methylation, and loading of the respective writer enzymes SUV39H1 and DNMT1. This dynamic process demonstrated a gradual transition from a transcriptionally active state, designated as the A-compartment to the inactive state of the B compartment altering the 3D epigenome. This is particularly of interest since Decitabine-induced genome-wide DNA hypomethylation resulted in large-scale 3D epigenome deregulation, including decompaction of higher-order chromatin structure and loss of boundary insulation of topologically associated domains (Achinger-Kaweck et al., 2024). The authors also demonstrated DNA hypomethylation resulted in ectopic activation of ER-enhancers, a gain in ER binding, the creation of new 3D enhancer–promoter interactions, and concordant up-regulation of ER-mediated transcription pathways. This phenomenon was reversible, since long-term withdrawal of it partially restored DNA methylation at ER-enhancer elements and loss of ectopic 3D enhancer–promoter interactions along with gene repression (Achinger-Kaweck et al., 2024). Therefore, targeting the 3D epigenome may open avenues for cancer therapy.

In summary, here we demonstrated a novel epigenetic mechanism of RNA Pol I inhibitor CX-5461 in mammalian cells. Gradual disruption of nucleolus structure and NPM1 localization from nucleolus to LADs led to epigenetic landscape changes with orchestration of heterochromatin state. Despite the lack of ribosome biogenesis, the enzymatic steps of the eraser and writer enzyme ensured remodeling of the 3D epigenome. Therefore, RNA Pol I mediated transcription may be associated with 3D genome and epigenome architecture.

## Materials and Methods

### Chemicals and cell exposure

CX-5461 and BMH-21 were purchased from Millipore-Sigma (# 5092650001, # SML1183 and # 58880-19-6, respectively). CX-5461 and BMH-21 were used at 10 mM.

### Immunofluorescence

HT1080 cells were grown according to ATCC’s recommendations on slides (VWR micro cover glass # 48366067) in 6 well plates to 80% confluency with or without 10 mM of CX-5461 for different time points. Cells were crosslinked with 4% formaldehyde for 10 min and quenched with 1.5 M Tris.Cl pH 8 for 5 min. In some experiments, cells were washed once with 1X PBS before crosslinking and 700 ml of CSK buffer was added for 5 min at 4°C. 500 ml of 1X PBS + 5 mM MgCl_2_ were added for 45 min at 37°C with or without RNase A (NEB # T3018-2) or RNase H (NEB # M0297S). After formaldehyde quenching, cells were permeabilized with 100% MeOH for 20 min at −20°C and incubated for 1h at RT with 5% BSA + 1X PBS + 0.3% Triton X-100. NMP1 Ab (Invitrogen # 32-5200, dilution 1/200) was incubated for 16 hr at 4°C. After 3 washes with 1X PBS + 0.1% Tween 20 for 5 min at RT, anti-lamin B2 (Invitrogen # PA5-29121) was added (1/200 dilution in 5% BSA + PBS + 1X + 0.3% Triton X-100) and incubated for 16 hr at 4°°C. NPM1 was revealed with anti-mouse IgG Alexa 594 (Invitrogen # A-11032) and lamin B2 was detected with an anti-rabbit IgG Alexa 488 (Invitrogen # A-11008). After 3 washes with 1X PBS + 0.1% Tween 20 (5 min at RT), slides were dried and mounted using Prolong Gold antifade reagent with DAPI (Invitrogen # P36931). Immunofluorescent images were acquired using Zeiss confocal LSM 880 microscope.

### EM-Seq

200 ng of HT1080 genomic DNA was used for detecting methylated cytosine. The protocol was performed according to manufacturer’s recommendations (NEB # E7120S).

### NEED-seq

NEED-seq size selection protocol was used as described previously (Sen et al., 2025). Briefly, HT1080 cells were grown on 6 well plates to 80% confluency and incubated with 10 mM of CX-5461 for different times. Cells were then incubated for 5 min at 4°C using CSK buffer to remove cytoplasm, followed by crosslinking with 4% formaldehyde for 10 min at RT. 1.5 M Tris.Cl, pH 7.5 was used to quench formaldehyde for 5 min at RT. After 1X PBS wash, blocking buffer was added to cells for 1 hr at RT. Different antibodies (see Table) were then added at dilutions recommended by the manufacturer and incubated with cells for overnight at 4°C. After 3 washes with 1X T-PBS for 5 min at RT, blocking buffer was added with fluorophore conjugated antibody for 1 hr at RT. After 3 washes with 1X T-PBS for 5 min at RT, NEED-seq binding buffer was added along with Nt.CviPII-pGL enzyme for 1h at RT. After 3 washes with NEED-seq washing buffer for 15 min at RT, 1X NEBuffer™2 was added for 1 hr at 37°C allowing frequent nicking activity of Nt.CviPII-pGL to fragment the targeted DNA. After 1 wash with 1X PBS, genomic DNA extraction was performed using Monarch Genomic DNA purification Kit (NEB # T3010S) with a minor modification which included a longer lysis step at 56°C for 2 hr to reverse DNA crosslinking. 1 µg of genomic DNA was used for size selection using first 0.4X volume of NEBNext Sample Purification beads (NEB # E7767S) with 10 min incubation at RT. Supernatant containing 200-500 bp DNA sizes was separated from beads using 12-tube magnetic separation rack (NEB # S1509S) and collected. An additional 0.4X volume of NEBNext Sample Purification beads was added to the supernatant and incubated for 10 min at RT. After 2 washes with 80% EtOH, DNA was eluted from beads with 50 µl of low TE buffer and DNA libraries were made using NEBNext Ultra II DNA Library Prep (NEB # E7103L) with some modifications. Briefly, end repair/dA tailing was performed on size selected DNA and NEBNext Illumina adaptor (1/10 dilution) were ligated for 1h at RT. USER enzyme (NEB # E7103L) was incubated for 20 min at 37°C and another round of library size selection was performed as described above. The library was amplified using 12 PCR cycles. PCR products were purified using 0.9 X volume of NEBNext Sample Purification beads. 1 nM of size selected DNA library was sequenced on Novaseq 6000. The list of antibodies is given in Suppl Table 1.

### NicE-seq

In the modified version of NicE-seq (Ponnaluri et al., 2017; Esteve et al 2020) HT1080 cells were grown on 6 well plates to 80% confluency and incubated with 10 mM of CX-5461 for 9hr. Cells were then incubated for 5 min at 4°C using CSK buffer to remove cytoplasm, followed by crosslinking with 4% formaldehyde for 10 min at RT. 1.5 M Tris.Cl, pH 7.5 was used to quench formaldehyde for 5 min at RT. After 1X PBS wash, 1 ml of Nt.CviPII (NEB # R0626S) was added per well in 800 ml of NEBuffer™2 1x for 1h at 37°C. Reaction was stopped with 5 mM of EDTA. After one wash with 1X PBS, genomic DNA was purified, size selected, and libraries were made using NEED-seq protocol (see above).

### Hi-C library preparation

HT1080 cells (0.5×10^6^) were incubated with or without 10 mM of CX-5461 for 9 hr. Hi-C libraries were prepared using protocol described previously (in Sen et al. 2025).

### RIP-seq

One 150 mm petri dishes of HT1080 cells used with or without 10 mM of CX5461 for 9 hr. After trypsinization, cell pellet was washed once with 1X PBS (spin at 1000 rpm at RT). Pellet were resuspended on ice with CSK buffer for 5 min, spun at 2000 g for 5 min at 4°C. Supernatant was discarded and pellet washed once with CSK buffer and resuspended in 1 ml of 1X PBS with 1% formaldehyde for 10 min at RT. 250 ml of 1.25 M glycine was added to quench formaldehyde for 5 min at RT. Nuclei were spun at 2000 g for 5 min at 4°C and washed once with 1X PBS. Nuclei (pellet) resuspended in 200 ml of 50 mM Tris.Cl pH 7.5, 200 mM NaCl and 1% SDS at RT. 200 ml of 10% Triton X-100 was added to quench SDS and crosslinked nuclei were sonicated 12 times (10 pulses each time). Debris (pellet) removed by 2 min 14000g spin at 4°C and supernatant kept for immunoprecipitation (IP). 200 mg of nuclear proteins used for IP (100 ml) were diluted in 900 ml of TD buffer (50 mM HEPES, pH 7.5, 250 mM NaCl and 1% Triton X-100). Precleaning was performed by adding 50 ml of protein G magnetic beads (NEB # S1430S) for 1h at 4°C. Supernatant was collected and 2.5 mg. of anti-mouse IgG (Santa Cruz # sc-2025) or anti-NPM1 (Invitrogen # 32-5200) antibody were added for IP overnight at 4°C. After IP, beads were washed 3 times (5 min at 4°C) with TD buffer and resuspended in 100 ml of 50 mM Tris.Cl pH 7.5, 200 mM NaCl, 1 % SDS including 2 ml of proteinase K (NEB # P8107S) overnight at 65°C, 1400 rpm. Input control is 0.37x. 12 ml of 10% Triton X-100 added to quench SDS. Supernatant was collected after separating beads on magnetic rack. 1 ml of TRIzol (Ambion) was added to supernatant for 5 min at RT and 200 ml of chloroform added per ml of TRIzol. After vortexing for 15 sec vigorously and incubation for 2-3 min at RT, samples were centrifuged for no more than 12,000 g for 15 min at 4°C. Aqueous phase was collected where RNA remains (2 mg of glycogen was added). The volume of aqueous phase (about 60% of Trizol volume) was measured and 0.5 ml isopropanol per ml of TRIzol added. Samples were then incubated at −80°C for 2 hr and centrifuged for no more than 12,000 g for 10 min at 4°C. RNA pellets were washed once with 75 % EtOH, dried and resuspended in RNAse free water (44 ml). DNase 1 reaction buffer (5 ml) and DNase 1 enzyme (NEB # M0303S) were added for 30 min at 37°C. 90 ml (1.8x) NEBNext RNA Sample Purification Beads (NEB # E7767S) to capture RNA and were incubated for 15 min on ice. Beads were washed twice with 80% EtOH and RNA was eluted with 6 ml of RNase free water. 5 ml of RNA was mixed with 4 ml of NEBNext First Strand Synthesis reaction buffer (lilac) and 1 ml of random primers (lilac) (reagents from NEBNext UltraII Directional RNA for Illumina; NEB # E7550S). Samples were incubated for 7 min at 94°C and were transferred immediately to ice. 8 ml of NEBNext Strand Specificity Reagent (brown) and 2 ml of NEBNext First Strand Synthesis Enzyme mix were added and incubated first at 25°C for 10 min, then 42°C for 15 min and 70°C for 15 min. After samples were put on ice, second strand cDNA synthesis was performed using 8 ml of NEBNext Second Strand cDNA Synthesis Reaction Buffer with dUTP mix (orange), 4 ml of NEBNext Second Strand cDNA Synthesis Enzyme Mix (orange) and 48 ml of nuclease free water for 1h at 16°C. 144 ml (1.8x) NEBNext RNA Sample Purification Beads (NEB # E7767S) were added and incubated for 5 min at RT. Beads were washed twice with 80% EtOH and double strand cDNA was eluted using 50 ml of 0.1x TE buffer. Samples were stored at −20°C until the next day. DNA repair/dA tailing was performed (UltraII protocol for Illumina; NEB # E7546S). Illumina adapters were diluted 1/5 for input; 1/25 for IP samples and ligated for 2hr at RT. USER enzyme was added (3 ml per sample; NEB # M5505S) for 20 min at 37°C. 87 ml (0.9x) of NEBNext RNA Sample Purification Beads was added for 10 min at RT. After 2 80% EtOH washes, elution was done with 16 ml of 0.1x TE buffer. 15 ml was used for PCR (14 cycles) with NEB # E6443A SET2 primers. After PCR, 45 ml (0.9x) of NEBNext Sample Purification Beads were added for 10 min at RT, washed twice with 80% EtOH. Elution was performed using 12 ml of 0.1x TE buffer.

### esiRNA knockdowns

20 nM of esiRNA against NPM1, RNA polymerase I, HDAC1, SUV39H1 or GFP (Sigma Aldrich # EHU115611 # EHU005201, EHU025841, EHU158181 or # EHUEGFP, respectively) were transfected using HiPerfect reagent (Qiagen #301705) in HT1080 cells for 24 hr. esiRNA GFP was used as control. After 3 successive transfections, cells were either fractionated (see below) or total protein cell content was extracted using 100 ml of 50 mM Tris.Cl pH 7.5, 200 mM NaCl and 1% SDS. 100 ml of 10% Triton X-100 was added to quench SDS. Samples were sonicated 10 times (10 pulses each time; Heat Systems, Ultrasonics, Model W-225R). Protein quantification with BCA kit. 2 mg total protein extracts or chromatin-bound or -unbound proteins were loaded on SDS-PAGE for Western blotting. The list of antibodies is given in Suppl Table 1.

### Cellular fractionation

Cytoplasm, chromatin bound, and unbound proteins were fractionated using Mayer and Churchman protocol (Mayer et al. 2017). Chromatin fraction was resuspended in 100 ml of 50 mM Tris.Cl pH 7.5, 200 mM NaCl and 1% SDS at RT. 100 ml of 10% Triton X-100 was added to quench SDS. Chromatin fractions were sonicated 10 times (10 pulses each time). Protein quantification with BCA kit. 2 mg of bound or unbound chromatin proteins was loaded on SDS-PAGE for Western blotting.

### HDAC1 and SUV39H1 assays

For HDAC1 assay, HeLa Core Histones (Active Motif # 53501) were incubated with different concentrations of recombinant NPM1 (Origene # TP303841) with or without recombinant HDAC1 (Abcam # 101661) in 20 ml reaction at 37°C for 4 hr. Blue Protein Loading Dye (NEB # B7703S) was added to the reaction. Samples were boiled and loaded on SDS-PAGE for NPM1 (CST # 92825S), HDAC1 (CST # 34589S), H3K9ac (CST # 9649S), H3K27ac (CST # 8173S) and histone H3 (CST # 4499S) Western blotting. The percentage of histone H3 acetylation was determined by densitometry using Image J.

Recombinant histone H3K9ac (EPL) (Active Motif # 31253) was incubated with different concentrations of recombinant NPM1 with or without recombinant HDAC1 in 20 ml reaction at 37°C. After 4 hr reaction, 200 ml of TD buffer (50 mM HEPES, 250 mM NaCl and 1% Triton X-100) was added and recombinant histone H3K9ac was immunoprecipitated using 5 ml of anti-C-term histone H3 antibody (CST # 14269) during 1 h 4°C. 20 ml of protein G magnetic beads (NEB # S1430S) were added for an additional 1h at 4°C. After histone H3 capture, beads were washed twice with 200 ml of SUV39H1 buffer (50 mM Tris.Cl pH 8.6, 5 mM MgCl2, 4 mM DTT and 20 mM of S-adenosylmethionine, NEB # B9003S) using magnetic separation rack (NEB # S1506S) and were resuspended with 200 ml of SUV39H1 buffer. Recombinant MBP-SUV39H1 was added to allow H3K9me3 reaction for overnight at RT. Beads were then captured on magnetic rack, supernatant was removed. Beads were resuspended in Blue Protein Loading Dye) and boiled. Beads were separated using magnetic rack and supernatant was loaded on SDS-PAGE. For HDAC1 assay and subsequent SUV39H1 assay, Western blot was performed using anti-H3K9ac (CST # 9649S) and anti-H3K9me3 (CST # 13969S). Anti-histone H3 (CST # 4499S) was used as control. The percentage of histone H3 acetylation and methylation were determined by densitometry using Image J.

### NPM1 Diffusion calculation from Microscopic Images

We have used microscopic images for 0h to 15h with three color channels i.e., (1) DAPI for DNA labeling; (2) Fluorescein Green for NPM1, and (3) Texas Red for Lamin B2. To calculate diffusion of NPM1 over the temporal landscape, we have implemented a special application of Stokes-Einstein Theory:

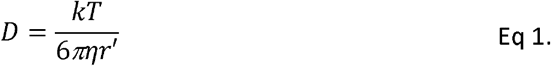

where,

*k* = Boltzmann constant

*η* = Viscosity

*T*= Temperature (in kelvin)

*r*’= hydrodynamic radius

By definition, *r*’ ∝ mass. In this case, we have considered the ratio of Intensity of fluorescence (I_NPM1_) and Intensity of DAPI (I_DAPI_), therefore, (I_NPM1_ / I_DAPI_) ≡ Relative mass. Therefore, the value of *r* is depending on 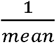 of relative mass.

Similarly, 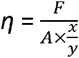 where,

*A* = Area of nucleus (ellipse like structure) = *π(r_1_+r_2_)* where *r*_*1*_ = shortest radius and r_*2*_ = longest radius

*F* = Intensity of NPM1

*x*/*y*= rate of shear Deformation ≡ Decay Rate *ln(k)*

We have considered 100 nuclei as input for each time points.

### NEED-seq and NicE-seq

NEED-seq and NicE-seq analysis was started from Adaptor trimming. Applying Trim Galore, low-quality sequences and adaptor sequences were trimmed (http://www.bioinformatics.babraham.ac.uk/projects/trim_galore) from paired-end sequencing reads using the commend line interface (CLI): --clip_R1 4 –clip_R2 4 –three_prime_clip_R1 4 – three_prime_clip_R2 4. Subsequently, Bowtie2 (Langmead et al., 2012) was applied on trimmed read pairs to map reference genome (hg38) with the following setup: --dovetail –no-unal –no-mixed –no-discordant –very-sensitive -I 0 -X 1000. After getting sam alignment files from Bowtie2, the files were converted to bam files using “samtools view -S -b” (Li et al., 2015) which were further purified by removing PCR duplicates using following settings: “picard MarkDuplicates REMOVE_DUPLICATES=true” and mitochondrial reads (http://broadinstitute.github.io/picard/).Resultant aligned read pairs were utilized for peak calling with MACS3 (Zhang et al., 2008). From NEED-seq, we generated two different set of peaks, (1) Macs3 call peak -f BAMPE -m 4 100 –bdg –SPMR (for narrow peak); (2) Macs3 call peak -f BAMPE -m 4 100 –bdg --broad –SPMR (for broad peak). For NicE-seq, we have called only narrowPeaks. We have generated three replicates for NicE-seq at Control as well as CX-5461 treated conditions. The analysis steps are combined in a nextflow pipeline.

### Methylation data processing

EM-seq methylation data was used for CpG methylation study. Quality check and removal of the low-quality sequences were performed following the same pipeline that was used in NEED-seq. Further, the fastq files were aligned with reference genome (hg38) using bwameth.py (https://github.com/brentp/bwa-meth). PCR duplicates were removed using “samblaster -e -i” (Faust et al., 2014). Next, mitochondrial reads were removed. Resultant bam files were utilized for further analysis. The methylation call was performed using MethylaDackel extract (github.com/dpryan79/MethylDackel), with coverage 25X threshold and more than 25% per CpG methylation.

### Hi-C data processing and upstream analysis

For Hi-C data processing, HiC-Pro (V.3.1.0) pipeline was utilized (Servant et al., 2015). HiC-Pro has five processing stages and one quality check stage. For data processing, we have considered MAPQ ≥ 30, Mbo I ligation sites (GATC) for resolution 50kb and 25kb for hg38 assembly. For ICE normalization, MAX_ITER was provided as 100. Control condition has seven pair-end replicates whereas treated condition has five pair-end replicates. After generating aligned bam files considering MAPQ threshold, we have merged replicates and performed the subsequent stages for generating “allValidPairs”. To visualize the TADs, we have also utilized “hicPlotTADs”.

### RIP-seq data processing

For RIP-seq data processing, we have followed the NEED-seq/NicE-seq pipeline. However, we have used T2T (V2) assembly in place hg38 assembly for NPM1 bound RIP-seq data. By the definition, T2T assembly is well characterized for non-coding rRNA. Therefore, we have it to ensure less noisy data collection. We have also included IgG and Input.

### Generating count matrix and noise reduction

As mentioned earlier, handling RIP-seq noise especially for non-coding RNA is critical. First, we called the count matrices based on T2T gtf annotation files from UCSC genome browser for NPM1, Input, and IgG. Now, input RIP-seq can feature the total count value for each RNA where IgG RIP-seq can show random targeting. Therefore, we start with normalizing data using count per million (CPM). Next, NPM1 count matrices was corrected based on IgG RIP-seq. Finally, we assessed absolute count based on percentage of count, comparing to Input.

### Differential peak analysis

To identify differential accessible chromatin peaks, DiffBind (Stark et al. 2011), a R package, was utilized. We have generated three replicates each for controlled and treated condition. Generated peak for each replicated along with the bam files were provided as input. DEseq2 option was selected in dba.analyze command where peaks with p-value ≤ 0.05 were considered as differential peak. Next, volcano plot was generated using dba.plotVolcano where log_2_FC ≤ −1 were considered as −ve enriched peaks and vice versa.

### Correlation and enrichment analysis

Biological replicates were merged using “samtools merge” (Li et al., 2015). Subsequently, merged bam files were downsized to the same range using Sambamba (Tarasov et al., 2015). Further, the peaks were generated using MASC3 settings as described before (Zhang et al., 2008). We followed pipeline suggested in Sen et al. 2025 for correlation analysis. Next, deeptools plotCorrelation (--whatToPlot heatmap) was used for calculating and plotting the correlation coefficient value. Enrichment analysis needs bigwig files. Initially, all the bam files were converted to bigwig files following “bamCoverage – normalizeUsing RPKM” where reads were normalized applying RPKM measure. These bigwig files can further be utilized for signal-to-noise visualization using Integrative Genome Browser (IGV) tools (Robinson et al., 2011). Further, matrices for enrichment were calculated using “deeptools computeMatrix” for both “reference-point” and “scale-regions” modes considering respective peak regions and TSS regions for hg38 downloaded from UCSC genome browser using the generated bigwig files. The resultant matrices were used further as input for “plotProfile” and “plotHeatmap” (Ramírez et al., 2010). Mostly, enrichment plots were generated for ± 2Kb up-stream and down-stream for NEED-seq/NicE-seq peaks except for LAD domains and Hi-C compartments where we considered ± 20Kb and ± 5Mb up-stream and down-stream respectively. Next, we utilize “bioframe” package (open2C et al. 2024) was used to summarize NEED-seq bigwigs in 25kb bins and generate pandas dataframe. Further, seaborn KDEplot were performed to generate KDE plots.

### Identification of LAD domains

IgG bam files were utilized for identification of lamin B2 LAD domains using bamCompare Command Line Interface (CLI): --scaleFactorsMethod SES – operation log2ratio -bs 20000 (Ramirez et al., 2010). Positive log2 values were considered as LAD domains.

### Identify Nucleolar Associated Domains (NADs)

NADfinder (Vertii et al. 2019), a R package, was used to predict Type I NADs, Type II NADs, and NPM1 NADs from H3K9me3, H3K27me3 and NPM1 NEED-seq bam files using IgG as background control. While calling the domains, we have considered high stringent padj value ≤ 0.001.

### Bed regions overlap

While analyzing the data, we have utilized three types of bed regions, 1. Peaks file, 2. Domain files, and 3. Methylation file. For demonstrating converagence of first two types, overlapping bed region were defined using “intervene venn” (Khan et al., 2017) where we utlized a parametric options “--bedtools-options f,r=0.5”. This parameter defined that at least 50% overlap from both bed files were considered as true convergence. As domains were mostly large in size, this stringent cutoff was essential to maintain robustness. Similarly, methylation beta value distribution in peak regions were identified using “bedtools intersect”.

### Subcompartments analysis

Subcompartments were based on Hi-C and NEED-seq Lamin B2 and NPM1 data. We confine the Hi-C and Lamin B2, NPM1 with fixed range by using principal component 1 (PC1) from Hi-C (generated using runHiCpca from HOMER; Heinz et al., 2010) and Lamin B2 and NPM1 genome wide domains averaged to 25 kb bins using deeptools MultiBigwigSummary exclusively. Further K-means clustering (K=10) was performed on summarized files separately. The resultant clusters were termed as subcompartments for Lamin B2 and NPM1 (Siegenfeld et al., 2022). Next, heatmaps were genarated between Lamin B2 and NPM1 clusters, using the Z-normalize Jaccard coefficient (Sen et al. 2025).

### A/B subcompartments and differential loops

“.allValidPairs”, resultant files from HiCPro, were utilized to calculate differential loops and A/B subcompartments. For A/B subcompartments,.allValidPairs were first converted to.hic files using hicpro2juicebox.sh. These.hic files were further used as input in Calder (Liu et al. 2021), a R package where bin size was given as 25Kb and assembly was hg38. Similarly, cLoops2 (Cao et al. 2022), a python API was implemented for differential loops calculation. This process consists of few sequential stages. First, “.allValidPairs” files were converted “.bedpe” files. Next, “cLoops2 pre” were performed to generate chromosome wise cis interacting “.ixy” which were further used as input for “cLoops2 callLoops”. The resultant loop files for controlled and treated conditions were further used to determine differential loops applying “cLoops2 callDiffLoops”.

### Statistical testing

We have performed statistical testing on the densitometry from western blot and RIP-seq absolute count values between control and treated data. Two sample Student t-test was applied where p-value ≤ 0.05 was as significant. For the bar plot, we have considered mean ± SEM in Prism V.10.

## Figure Legends

**Supplementary Figure 1. RNA Pol I inhibition modulates NPM1 localization. A.** IGV screenshots of RNA Pol I and NPM1 binding on DNA using NEED-seq is shown. CX-5461 exposed conditions show loss of occupancy for RNA Pol I and NPM1 at the rDNA cluster. **B.** Comparative measurement of Diffusion Coefficient (DC), decay rate (ln(k)) and relative energy for NPM1 at various time interval post CX-5461 exposure is shown at various time interval of CX-5461 exposure. **C.** LOWESS smoothing plot exhibiting relation between relative energy and NPM1’s diffusion towards the nuclear periphery. **D.** LOWESS smoothing plot showing lamin signal re-distribution within nucleus at various time interval at various time interval of CX-5461 exposure. **E.** Western blot of NPM1 from control, and CX-5461 exposed cells (treated with RNase H) is shown. The corresponding bar plot (control in red, and CX-5461 exposed in green) is shown at the bottom panel. **F.** Western blots of total cell extract for RNA Pol I, lamin B2, and NPM1 at various time interval post CX-5461 exposure. Corresponding line graph demonstrates gradual decrease for RNA Pol I compared to at various time points as indicated post CX-5461 exposure.

**Supplementary Figure 2. NEED-seq of histone modifications of control and HT1080 cells exposed to CX-5461. A.** IGV screenshots showing NEED-seq on euchromatic (top) and heterochromatic (bottom) marks, as indicated on the left, along with control and CX-5461 exposed panels. **B.** Heatmaps based on Pearson’s correlation coefficient using NEED-seq reads to determine correlation with NPM1 and euchromatic marks as indicated for control and CX-5461 exposed cells. **C.** Heatmaps based on Pearson’s correlation coefficient using NEED-seq reads to determine correlation with NPM1, lamin B2 with euchromatic and heterochromatic marks in both control and CX-5461 treated cells.

**Supplementary Figure 3.Effect of RNA Pol I inhibitors on other histone modification. A.** Western blots of HT1080 cell extract (incubated with BMH-21, another RNA Pol I inhibitor) and corresponding bar plots, showing level of RNA Pol I, H3K9ac, and H3K9me3 along with p-values († p-value > 0.05; * 0.01< p-value ≤ 0.05; ** 0.001< p-value ≤ 0.01; ***p-value ≤ 0.001). **B-C**. Metagene plots (left panel), western blots (middle panel), and quantitative estimation in bar plot (right panel) are shown for H3K4me3 (B), and H3K27me3 (C) upon CX-5461 exposure. **D**. Western blots of chromatin bound (lower panel) and ch-unbound (nucleoplasm, upper panel) fraction (left panel) and corresponding relative quantity of RNA Pol I with histone H3 and NUP153 (right) are shown post CX-5461 exposure at various time interval as indicated. **E-F**. KDE plots demonstrating NPM1, H3K9ac and H3K9me3 coverage at 0 and 9hr post CX-5461 exposure. Please note NPM1 (green fill) shifts upon 9hr CX-5461 incubation contributing to loss of H3K9ac density (red line plot, left panel) and re-distribution of H3K9me3 (blueline plot, right panel), compared to NPM1 (red fill plot), H3K9ac (green line plot, left panel), and H3K9me3 (yellow line plot, right panel) at 0h, demonstrating NPM1 re-distribution in chromatin landscape.

**Supplementary Figure 4. siRNA mediated knockdowns and their effect on histone modifications. A.** Western blots and associated bar graphs following siGFP control (sea green) and 3x siPol I (pink triangles) transfection into HT1080 cells. Upper panel shows the level of RNA Pol I, HDAC1, H3K9me3, and NPM1 in chromatin bound fraction in western blot (left panel). These proteins were normalized to histone H3 and represented as bar graphs (right panel). Middle panel same transfection as the upper panel with H3K9me3, H3K27me3 and H3K4me3 western blot (at the left) and their quantification normalized to histone H3 (at the right) is shown. Lower panel same transfection as the middle panel with RNA pol I, HDAC1, and NPM1 western blot (at the left) and their quantification normalized to NUP153 (at the right) is shown. Increase or decrease is shown with p-values († p-value > 0.05; * 0.01< p-value ≤ 0.05; ** 0.001< p-value ≤ 0.01; ***p-value ≤ 0.001). **B.** Comparative western blots and associated on H3K9me3 from total cellular extract using siHDAC1 and siSUV39H1 transfection along with siGFP in control and CX-5461 exposed cells, showing the impact on total HDAC1 and SUV39H1. Quantity of proteins from western blot are normalized to histone H3 and corresponding bar graphs are shown in the right panel. **C.** Western blots (top panel) and corresponding bar graphs (bottom panels) for H3K9ac (dark pink) and HDAC1 (green) total extract upon control and CX-5461 exposure normalized to histone H3 is shown.

**Supplementary Figure 5. Western blots on chromatin bound fraction of DNMTs A.** At various time interval post CX-5461 exposure to HT1080 cells **B.** Corresponding line graph of chromatin bound DNMTs. Note, gradual increase for DNMT1 on chromatin compared to DNMT3A/3B at various time points as indicated post CX-5461 exposure (normalized to Histone H3).

**Supplementary Figure 6. Chromatin state alteration and 3D changes post RNA Pol I inhibition. A.** ChromHMM state showing a global transition of NAD I, NAD II and LAD regions. NAD I and LAD regions are distributed within TSS, genic enhance regions and transcriptionally active regions. NAD II are lost at 5’ and 3’ regions of genes along with genes associated with strong transcription upon RNA Pol I inhibition, as shown. **B.** Z-normalized Jaccard coefficients using heatmap between NPM1 and LAD subcompartments (generated by performing K-means (K=10) clustering on summarized PC1 (Hi-C) and NEED-seq signals within 25 kb bins), are presented showing NPM1 distributions within LAD regions in 3D context in control (left panel) and following RNA Pol I inhibition (right panel). **C.** Bar graph showing the number of TADs between control and CX-5461 exposed cells. **D.** Box plots showing the change in TAD size in control and CX-5461 treated cells. **E.** Bar graphs showing a significant shift of TADs within LAD region in CX-5461 treated cells compared to control cells.

**Supplementary Figure 7. TAD remodeling and NPM1 redistribution in A and B compartments. A.** Hi-C plots from control and CX-5461 exposed cells showing TADs domain expansion within LADs in chr 21. **B.** Metagene plots showing gradual NPM1 enrichment from A- to B-compartment subtypes regions (± 5 Mb) in CX-5461 exposed (green) and control (red) cells.

**Supplementary Figure 8. Genome wide loss of anchors and loops facilitating the heterochromatin spreading. A.** Genome Browser screenshot (from Wash U) showing significant loss of Hi-C loops corresponding to loss of NPM1 binding and H3K9ac occupancy (NEED-seq) signal upon RNA Pol I inhibition. **B.** Bar graphs exhibit ∼60% of the Hi-C loops differentially modified (I) among them ∼60% are negatively enriched (II). Majority (∼80%) of this negatively enriched HiC-loops are completely depleted upon RNA Pol I inhibition (III). A model of the depletion process is shown post CX-5461 exposure (IV). **C.** Metagene plots (± 2 kb) in anchors of depleted loop demonstrate loss of enrichment for NPM1 and H3K9ac with gain of H3K9me3 enrichment upon CX-5461 exposure.

## Data availability

NEED-seq, NicE-seq, RIP-seq, Hi-C and EM-seq data performed in this study are available in NCBI Gene Expression Omnibus (GEO).

## Acknowledgments

We thank T. Evans, D. Comb, Sir R.J. Roberts, and S. Russello for encouragement. The project was funded by basic research grant to SP from New England Biolabs, Inc., and partly funded by R44HG011875 from NIH.

## Declaration of generative AI and AI-assisted technologies in the writing process

During the preparation of this work the author(s) didn’t use any Generative AI assisted technology.

## Author contribution

SS, POE, RK performed experiments. Bioinformatic analysis was performed by SS. SP and AU conceptualized and planned experiments and wrote the manuscript with input from SS, POE and RK.

---

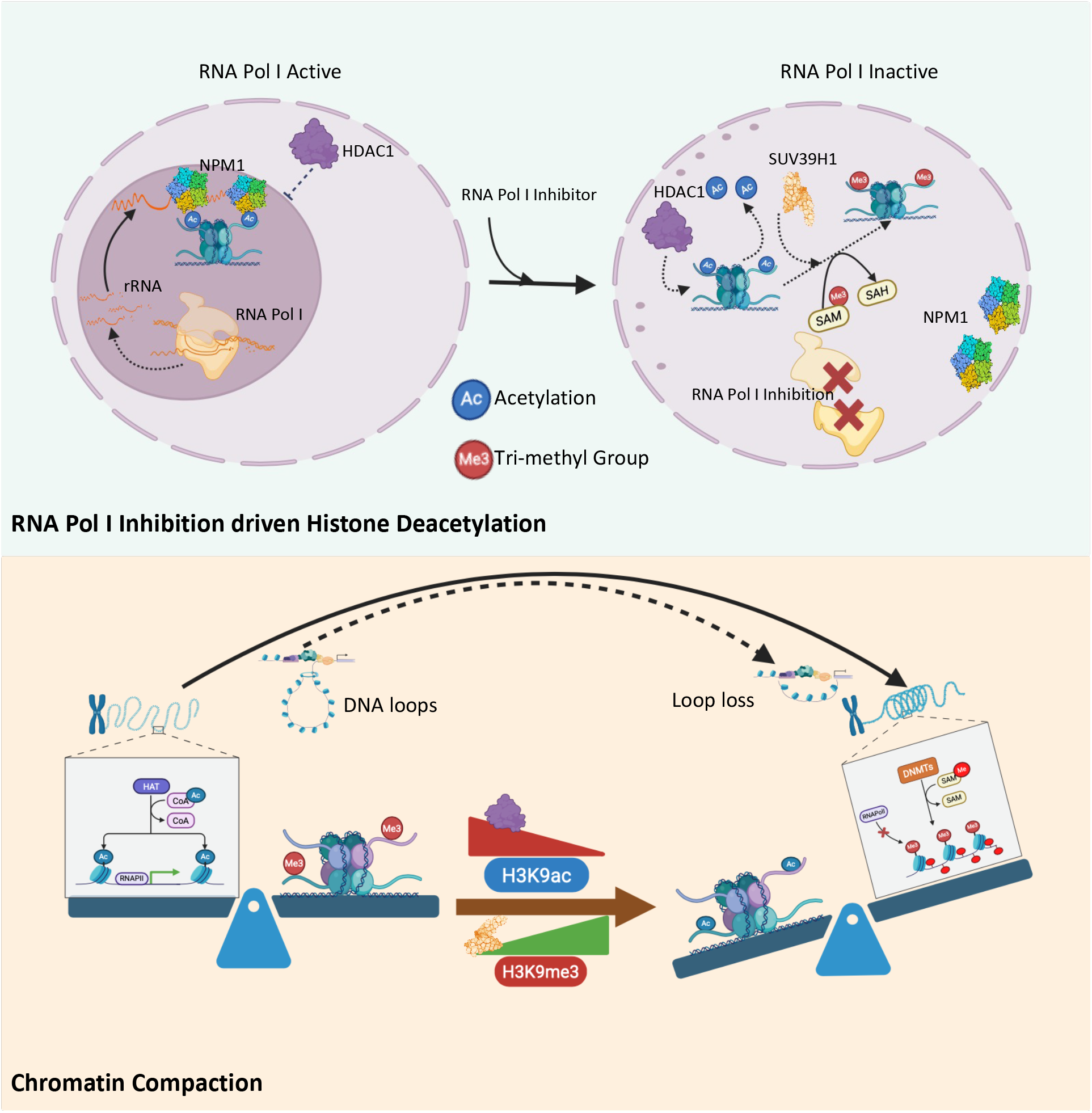

